# Spatial Gradient of Microstructural Changes in Normal-Appearing White Matter in Tracts Affected by White Matter Hyperintensities in Older Age

**DOI:** 10.1101/412064

**Authors:** Susana Muñoz Maniega, Rozanna Meijboom, Francesca M. Chappell, Maria C. Valdés Hernández, John M. Starr, Mark E. Bastin, Ian J. Deary, Joanna M. Wardlaw

**Affiliations:** Neuroimaging Sciences, Centre for Clinical Brain Sciences, University of Edinburgh, UK; UK Dementia Research Institute at the University of Edinburgh; Alzheimer Scotland Dementia Research Centre, University of Edinburgh, Edinburgh, UK; Department of Radiology and Nuclear Medicine, Erasmus MC – University Medical Centre Rotterdam, NL; Department of Psychology, University of Edinburgh, UK; Centre for Cognitive Ageing and Cognitive Epidemiology, University of Edinburgh, UK

**Keywords:** brain, ageing, diffusion MRI, white matter hyperintensities, tractography

## Abstract

Brain white matter hyperintensities (WMH), common in older adults, may contribute to cortical disconnection and cognitive dysfunction. The presence of WMH within white matter (WM) tracts indicates underlying microstructural WM changes that may also affect the normal-appearing WM (NAWM) of a tract. We performed an exploratory study using diffusion magnetic resonance imaging of 52 healthy participants from the Lothian Birth Cohort 1936 (age 72.2 ± 0.7 years) selected to include a range of WMH burden, to quantify microstructural changes of tracts intersecting WMH. We reconstructed tracts using automated tractography and identified intersections with WMH. Tissue volumes and water diffusion tensor parameters (mean diffusivity (MD) and fractional anisotropy (FA)) were established for tract-WMH and tract-NAWM. MD and FA were also measured for tract-NAWM at 2 mm incremental distances from the tract-WMH edge, and from the edge of nearby, non-intersecting, WMH. We observed microstructural changes in tract-WMH suggestive of tissue damage. Tract-NAWM also showed a spatial gradient of FA and MD abnormalities, which diminished with distance from the tract-WMH. Nearby WMH lesions, not directly crossed by the tract, also affected tract microstructure with a similar pattern. Additionally, both FA and MD changes in tract-NAWM were predicted by FA and MD changes respectively in tract-WMH. FA was also predicted by tract-WMH overlap volume, whereas MD was better predicted by whole-brain WMH load. These results suggest that tract-NAWM microstructure is affected by the pathological process underlying WMH, when WMH are either within or near to the tract. The changes in NAWM tract tissue may indicate future lesion progression and may play an important role in cognitive ageing.

## 1. Introduction

White matter hyperintensities (WMH), or leukoaraiosis, is a condition routinely found in brain magnetic resonance imaging (MRI) scans of older people and contributes to cognitive and functional decline (Fernández-Cabello et al., 2016; Haynes et al., 2017; Ritchie et al., 2015b). These lesions appear as hyperintensities in the white matter (WM) on T2-weighted (T2W) or fluid attenuated inversion recovery (FLAIR) MRI, and have been described as WM degeneration characterized by axonal loss, demyelination and gliosis, on neuropathological examination (Fazekas et al., 1993).

It has been suggested that cortical “disconnection” due to the loss of WM fibres may play an important role in age-related cognitive decline (O’Sullivan et al., 2001; Ritchie et al., 2015a), and this disconnection could be the consequence of WMH within WM tracts. In support of this, several studies observed associations between WMH volume within specific WM tracts and cognitive performance (Biesbroek, 2016; Reginold et al., 2015) and functional connectivity (Langen et al., 2017; Taylor et al., 2017). These studies suggest that WMH may directly damage tracts and understanding the influence of WMH on WM tracts is therefore important in investigating age-related cognitive decline. This can be further investigated by assessing, not just the extent of the damage in the tract, but also the microstructural properties of WM tracts that are affected by WMH.

Microstructural qualities of WM can be assessed in terms of the water diffusion properties within the tissue, which is measured using diffusion MRI (dMRI), a quantitative MRI technique. Two widely used water diffusion biomarkers are fractional anisotropy (FA) and mean diffusivity (MD), obtained from the eigenvalues of the water self-diffusion tensor (Basser and Pierpaoli, 1996). FA is an indicator of the degree of directionality of water diffusion, and MD of the free water diffusion that can take place within the tissue. Both parameters are therefore sensitive to the underlying tissue microstructure, and can be used as probes to assess changes in structural barriers within WM, such as axonal membranes or myelin (Beaulieu, 2002). This has been supported by histology studies of WM which show that reduced axonal density, and anomalies in the myelin sheaths underlie abnormalities in these measures (Rodríguez-Cruces and Concha, 2015; Song et al., 2003).

Both FA and MD have shown age-related changes in WM tracts (Hsu et al., 2010; Mayo et al., 2017; McWhinney et al., 2016; Thomason and Thompson, 2011). These changes have been related to the ratio of overlap between WM tracts and WMH (for the remainder of the article referred to as *tract-WMH*), independent of the overall WMH burden (Maillard et al., 2015). This suggests that within-tract microstructural changes depend on the volume of tract-WMH and that individual tracts are affected to different levels. Additionally, FA and MD changes have also been observed in whole-brain normal-appearing white matter (NAWM)– i.e. abnormalities not yet visible–in the presence of WMH. This was particularly seen in NAWM areas surrounding WMH, with a diminishing effect of WMH on NAWM with distance from the visible WMH boundary (Maillard et al., 2011; Muñoz Maniega et al., 2015). This suggests that a similar distance pattern of NAWM tissue damage would accrue within specific tracts, which could be caused by the WMH within that tract, although they could also be caused by nearby WMH outside the tract. NAWM integrity changes have been shown to be more closely related with cognitive changes than the visible WMH (Schmidt et al., 2010; Vernooij et al., 2009). Therefore, in order to improve our understanding of age-related cognitive decline, the degree of disruption within tracts in not only visible WMH, but also in the NAWM, should be explored.

Since age has strong effects on white matter DTI parameters (Cox et al., 2016), in this study we used data from the Lothian Birth Cohort 1936 (LBC1936) a cohort of older age community-dwelling subjects, all born in 1936, with a very narrow age range at imaging, to minimise the effects of age. We investigate how WMH affect tracts and surrounding NAWM in older age, specifically, we hypothesise that tract-WMH will have a negative effect on the tissue integrity of the remainder, normal-looking WM of the tract (which will be referred to as *tract-NAWM*), and that this effect will become less apparent along the WM tract and away from the visible WMH. First, we assess the effect of tract-WMH load on water diffusion metrics within tract-NAWM. Second, we evaluate the water diffusion differences between tract-WMH and whole tract-NAWM. Third, we investigate water diffusion differences between tract-WMH and tract-NAWM at several distances from the tract-WMH in order to establish the extent of invisible damage in tract-NAWM. Lastly, we consider the potential influence of nearby WMH, with which the tract does not intersect, on tract-NAWM integrity to check whether nearby WMH affect tract-NAWM to the same extent as tract-WMH. As this study was exploratory to investigate proof of principle before application to the full cohort, we selected a subsample of the LBC1936 study, which represented the full range of WMH burden and without stroke.

## 2. Methods

### 2.1 Participants

The LBC1936 comprises a group of community-dwelling individuals born in 1936, most of whom took part in the Scottish Mental Survey of 1947. At approximately 70 years of age, the LBC1936 participants were recruited for follow-up cognitive and other medical and psycho-social assessments (Deary et al., 2012, 2007). During a second wave of this longitudinal study, at approximately 73 years of age, 700 participants underwent comprehensive MRI to assess brain structure (Wardlaw et al., 2011). Written informed consent was obtained from all participants under protocols approved by the National Health Service Ethics Committees.

The current study was an exploratory study using imaging data from the second wave of LBC1936. This pilot sample was chosen with three requirements: it represented all levels of WMH burden (Fazekas score (Fazekas et al., 1987)), each participant had structural and diffusion MRI data available, and participants did not have a history of stroke. We aimed to choose a sample of 60 participants, 10 with each Fazekas total score of 1 to 6 (sum of deep + periventricular 0-3 scores); however, only eight participants had a Fazekas score of 6 and without a stroke. We completed the sample with one extra case with each Fazekas score of 2 and 3, as these are the most frequent scores in the LBC1936. The final sample therefore consisted of ten participants with each Fazekas score of 1, 4 and 5; 11 participants with each Fazekas scores of 2 and 3; and eight participants with a Fazekas score of 6. We have showed previously strong correlation between Fazekas scores and WMH volume (Valdés Hernández et al., 2013b) therefore the chosen sample represented a full range of volumes in this cohort.

### 2.2 Imaging acquisition

MRI data were acquired using a GE Signa Horizon HDxt 1.5 T clinical scanner (General Electric, Milwaukee, WI, USA) using a self-shielding gradient set with maximum gradient of 33 mT/m and an 8-channel phased-array head coil. The full details of the imaging protocol can be found in (Wardlaw et al., 2011). Briefly, the MRI examination comprised a high-resolution 3D T1-weighted (T1W), T2W, T2*-weighted (T2*W) and FLAIR structural scans, as well as dMRI. The dMRI protocol consisted of seven T2W volumes (b = 0 s/mm^2^) and sets of diffusion-weighted (b = 1000 s/mm^2^) single-shot, spin-echo, echo-planar (EP) volumes acquired with diffusion gradients applied in 64 non-collinear directions (Jones et al., 2002). All sequences, except for the T1W, were acquired in the axial plane with a field-of-view (FOV) of 256 × 256 mm, contiguous slice locations, and image matrices and slice thicknesses designed to give 2 mm isotropic voxels for dMRI, and voxel dimensions of 1 × 1 × 2 mm for T2W and T2*W, and 1 × 1 × 4 mm for FLAIR. The high-resolution 3D T1W scan was acquired in the coronal plane with a FOV of 256 × 256 mm and voxel dimensions of 1 × 1 × 1.3 mm.

### 2.3 Visual scoring of white matter hyperintensities

WMH were defined according to the STRIVE criteria (Wardlaw et al., 2013). All assessments used validated visual or computational methods and were performed blind to all patient demographic, clinical and tractography characteristics. A qualitative assessment of WMH load was performed by an expert neuroradiologist who scored hyperintensities on the FLAIR and T2W scans using the Fazekas scale, after training on a standard data set. A total score ranging from 0 to 6 was obtained by summing the periventricular and deep WMH Fazekas scores. The Fazekas scale is one of the most widely used visual rating scales and has been in use for over two decades. To ensure observer reliability, a second consultant neuroradiologist cross-checked a random sample of 20% of ratings, all scans with stroke lesions, and any scans where the first rater was uncertain (Valdés Hernández et al., 2013b).

### 2.4 Whole brain WMH and NAWM segmentation

All structural MRI volumes were registered to the corresponding T2W volume using rigid body registration (Jenkinson and Smith, 2001). Whole brain NAWM and WMH tissue masks were obtained using the multispectral colouring modulation and variance identification (MCMxxxVI) method (Valdés Hernández et al., 2010). In brief, T2*W and FLAIR volumes were mapped into red-green colour space and fused; the minimum variance quantization clustering technique was then used in the resulting image to reduce the number of colour levels, thereby allowing WMH to be separated from other tissues in a reproducible and semiautomatic manner. The same method was used to extract the NAWM from the T1W and T2W volumes. Any stroke lesions (cortical, cerebellar, lacunes, and large subcortical) were identified by a neuroradiologist and excluded from the masks manually by a trained image analyst.

### 2.5 Diffusion tensor imaging analysis and tractography

dMRI volumes were pre-processed using FSL 4.1 (Jenkinson et al., 2012). First, brain extraction was performed using BET (Smith, 2002), and second, bulk motion and eddy current induced distortions were removed by registering all volumes to the first T2W EP volume (Jenkinson and Smith, 2001). Third, DTIFIT was used to obtain the water diffusion tensor on a voxel-wise level, and to calculate parametric maps of FA and MD from the diffusion tensor eigenvalues. This was followed by automatic tractography using Tracula implemented in Freesurfer5.3 (TRActs Constrained by UnderLying Anatomy; (Yendiki et al., 2011)). Tracula uses global probabilistic tractography (Behrens et al., 2007) and anatomical priors of the white matter pathways derived from a set of training subjects; its accuracy has been evaluated against manual tract labels (Yendiki et al., 2011). Registration to the tract atlas containing the priors was performed by affine registration to the MNI125 template (Jenkinson and Smith, 2001). We reconstructed the 18 white matter pathways included in Tracula (corpus callosum: forceps major and forceps minor, and bilateral corticospinal tract (CST), inferior longitudinal fasciculus (ILF), uncinate fasciculus (UNC), anterior thalamic radiation (ATR), cingulum: cingulate gyrus (CCG) and angular bundle (CAB), and superior longitudinal fasciculus: parietal (SLFp) and temporal (SLFt) segments). All white matter tracts were visually inspected and those not following the expected paths were discarded from further analysis. See Figure 1a and 1b for an example showing the 18 tracts and WMH in a representative participant.

**Figure 1.**
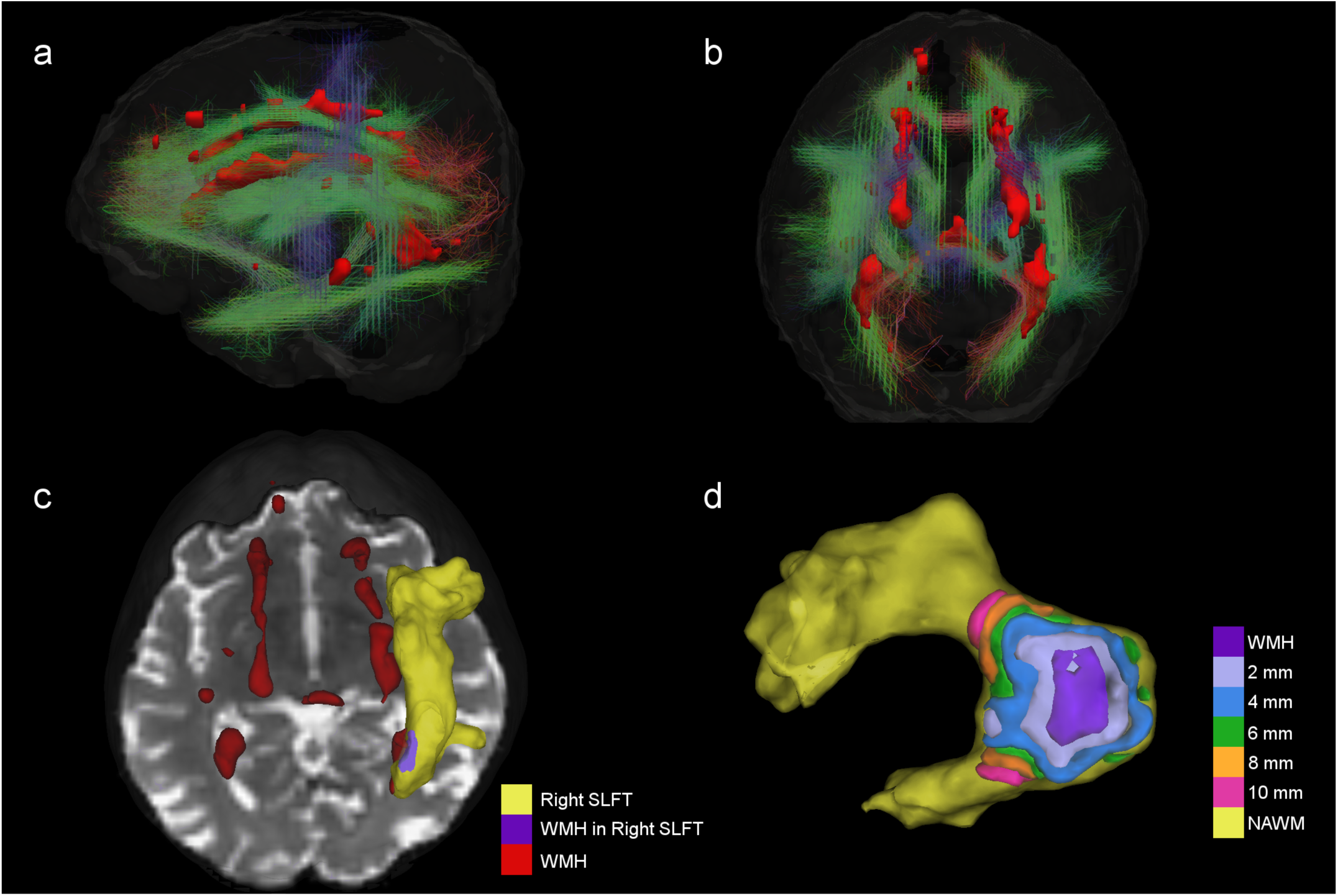
3D view of 18 tracts (streamlines) and WMH (red surface) from a participant with total Fazekas score of 3. a) and b) are the lateral and superior views of all tracts and WMH. Streamlines are colour coded according to the main direction of their middle segment of the tract (red = x axis, green = y axis and blue = z axis). c) example showing just the superior longitudinal fasciculus temporal (SLFt) ending as a yellow surface and the area of tract-WMH in purple, with remaining WMH in red. d) example of the contours used for the spatial analysis; the tract-WMH area (purple) is dilated by 2mm at a time to create surface contours within the tract-NAWM at different distances from the WMH edge.

### 2.6 WM tract-WMH and WM tract-NAWM intersections

To obtain the areas of the tracts that intersected with WMH for each individual, we first registered the T2W volume to the averaged T2W EP volume (S0) non-linearly using RNiftyReg (Clayden et al., 2015; Modat et al., 2010). This registration was then applied to the whole-brain WMH and NAWM masks created previously in order to overlap them with the Tracula tracts-masks in diffusion space and obtain the intersections. This way the tracts were divided into *tract-WMH* (as the intersection between the tract and the whole-brain WMH mask) and *tract-NAWM* (as the intersection between the tract and the whole-brain NAWM), see example in Figure 1c. These were subsequently overlaid onto the FA and MD parametric maps for quantitative measurements. Averages of FA and MD values from each tract area (WMH and NAWM) were obtained by weighting the corresponding voxels in the FA and MD maps by the posterior probability in those voxels given by the Tracula tractography outputs. Please note that not all tracts had a tract-WMH intersection, and the area of WMH overlap varied for different tracts and between participants. The percentages of overlap were quantified.

### 2.7 Spatial analysis of tracts

#### 2.7.1 Spatial contours

We assessed how the intersection or proximity of the WMH affected the tract integrity by creating approximately equidistant 3D contours around the tract-WMH, which propagated into the tract-NAWM for each tract. To achieve this, we dilated the tract-WMH masks by increments of 2 mm (1 voxel in dMRI-space) up to 10 mm, and then subtracted from each dilated ROI the previous ones. That is, the tract-WMH mask was subtracted from the 2mm ROI to obtain a contour at about 2mm from the WMH edge; the tract-WMH and 2mm ROI were subtracted from the 4mm ROI to obtain a contour at about 4mm from the WMH edge, and so on (the distances quoted are approximate as they are limited by the finite voxel size). For each contour, only the voxels overlapping with the tract-NAWM were kept for each tract, so no other tissues were included. See Figure 1d for an example of contours in 3D and ‘WMH1’ in Figure 2 for a graphical representation of this approach. Weighted means of FA and MD were obtained for each contour for parametric assessment of the effects of tract-WMH.

**Figure 2.**
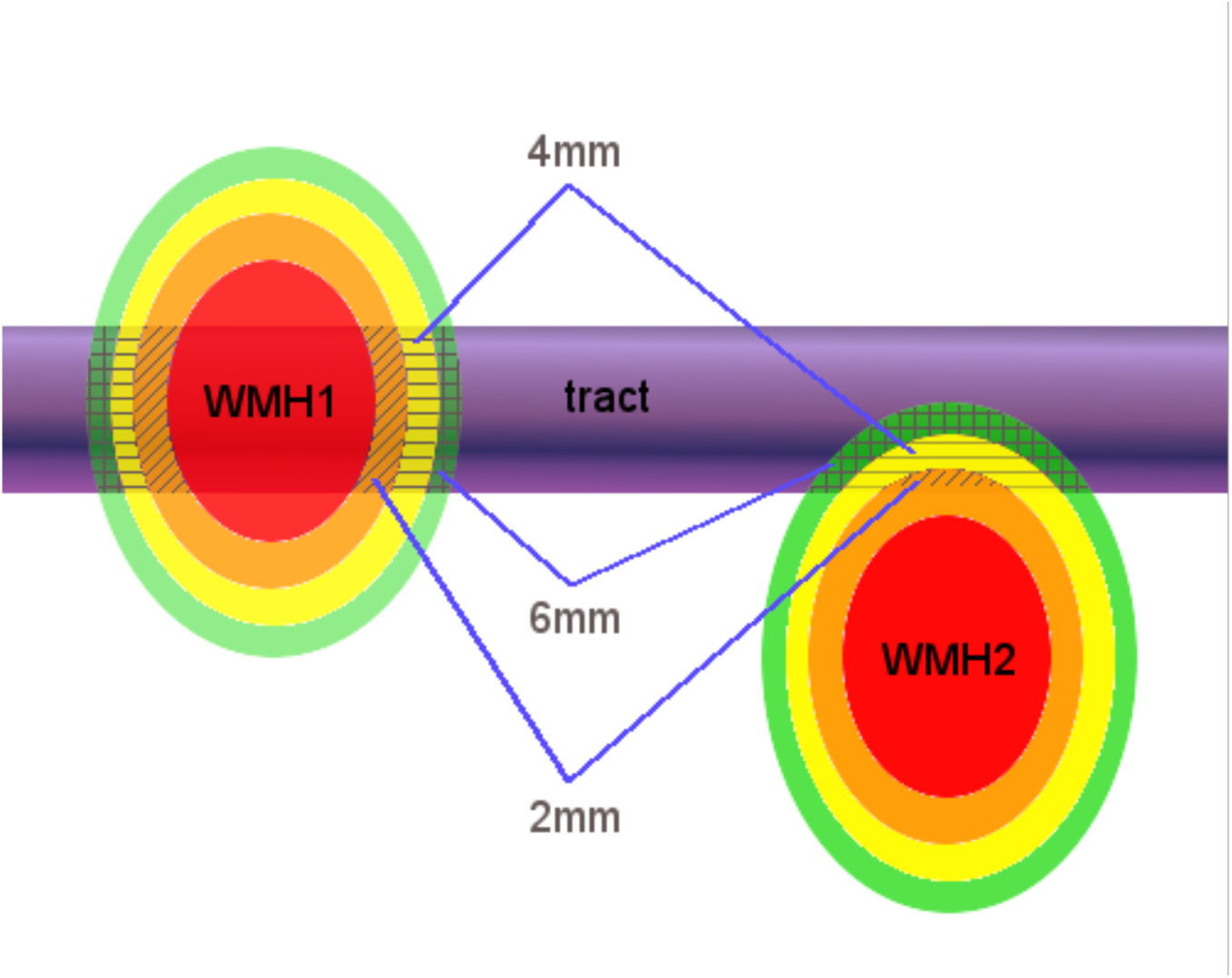
Schematic figure of the spatial analysis of the effects of tract-WMH (WMH1) and the nearby-WMH (WMH2) showing 2, 4 and 6mm contours around the WMH. Average FA and MD are measured on the intersections of the contours with the tract-NAMW only (patterned areas).

#### 2.7.2 Nearby-WMH

The spatial analysis was repeated for WMH that were nearby, but that did not intersect the tract, to assess their possible effect on the WM tract. We considered only those nearby-WMH for which any of the 2-10 mm contours intersected with the tract-NAWM (see ‘WMH2’ in Figure 2). Weighted means of FA and MD were obtained for each nearby-WMH contour that intersected with an individual tract-NAWM.

### 2.8 Statistical analyses

All statistical analyses were performed in R v.3.5 with packages *car* (Fox and Weisberg, 2011) and *lmerTest* (Kuznetsova et al., 2017). Plots were made with *ggplot2* (Wickham, 2016). Analyses were performed including all tracts within a model, and also for each individual tract, (results for individual tracts are provided in Supplement S2). For the models including all tracts, a second model was fitted including other potential confounders such as gender and age (number of days at time of scan).

Since the water diffusion parameters measured in tracts across the brain are highly correlated (Penke et al., 2010), a conservative Bonferroni approach for multiple comparison correction was not appropriate in the current analysis. However, to compromise between Type I and Type II error protection, we used a less conservative approach for correcting for multiple comparisons by setting the significance threshold at p < 0.01.

#### 2.8.1 Global and local WMH and tract-NAWM water diffusion

FA and MD values for tract-WMH and tract-NAWM were assessed for each individual WM tract and for all WM tracts combined, with measurements averaged for all tracts in each individual producing one value per participant. Additionally, the percentage of tract-WMH volume (% WMHvol) was established for each individual WM tract and for all WM tracts combined. These were calculated by dividing the tract-WMH volume by the total tract volume (tract-WMH plus tract-NAWM).

For all tracts combined, we investigated the predictive effect of tract-WMH diffusion (FA and MD respectively), mean tract % WMHvol, and Fazekas score on tract-NAWM FA and MD using a multiple linear regression analysis. The inclusion of these terms would provide an indication on whether NAWM integrity is better predicted by the extent of the localised damage within the tract (% WMHvol) or the global damage in the brain (Fazekas score).

A second multiple linear regression model was then run for both tract-NAWM FA and MD including gender and age. As an exploratory analysis, only the first regression model was applied to tract-NAWM FA and MD in individual WM tracts. A further analysis of water diffusion changes per Fazekas score is included in Supplement S1.

#### 2.8.2 Water diffusion in tract-WMH and tract-NAWM

A repeated measures linear mixed model with tissue type as a fixed effect was performed to compare FA and MD between tract-WMH and tract-NAWM. All tracts were included in the model by considering each tract measurement as a repeat. A model with random intercept and slope for both participant and tract provided the best fit (lowest Bayesian information criterion). A second model included age and gender to assess any significant effects. Type III Wald F tests with Kenward-Roger df approximation were used to obtain F and p-values. The analysis was repeated for each tract separately using a paired t-test.

#### 2.8.3 Water diffusion in tract-NAWM spatial contours for tract-WMH

We assessed the spatial changes of FA and MD values with distance from the tract-WMH with a repeated measurements linear mixed model of FA and MD measurements in tract-WMH, and in tract-NAWM at 2, 4, 6, 8 and 10 mm from the tract-WMH. As the trajectories of FA and MD show an asymptotic relationship of water diffusion with distance (Figure 5), log(distance+1) was used as fixed effect (with tract-WMH coded as 0 mm). All tracts were included in the model as repeats at each distance. A model with random intercept and slope for both participant and tract gave the best fit. Age and gender effects were included in model The analysis was repeated for each tract separately. Type III Wald F tests with Kenward-Roger df approximation were used to obtain F and p-values.

**Figure 5.**
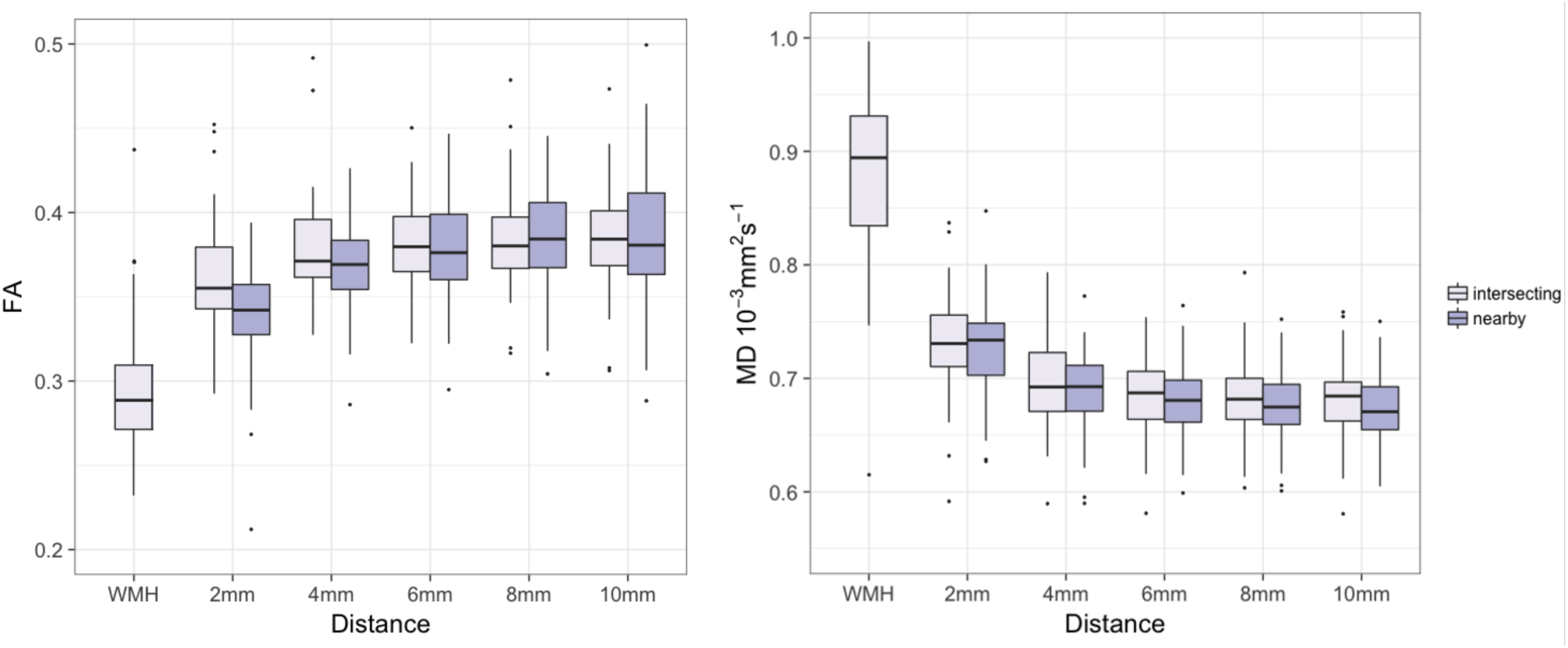
Water. **d**iffusion changes in tract-NAWM with distance to intersecting (tract-WMH) and nearby WMH. FA (left) and MD (right) averaged for all tracts. WMH measurements were only performed in tract-WMH. In all box plots, the boxes represent the lower and upper quartiles and the median measurement (thick line) for each group. Whiskers indicate the sample minimum and maximum, whereas the represented outliers (dots) differ from the lower and upper quartiles by more than 1.5 times the interquartile range.

#### 2.8.4 Water diffusion in tract-NAWM spatial contours for nearby-WMH

FA and MD values for spatial distances at 2, 4, 6, 8 and 10 mm from nearby-WMH were measured for all tracts. We then assessed the effect of the two types of WMH (tract or nearby) on the water diffusion measurements in NAWM spatial contours. The value of WMH was not available for nearby-WMH, hence we compare the trajectories between tract-WMH and nearby-WMH for distances 2-10 cm only. We used a repeated measures linear mixed model with distance and WMH type as fixed effects. The best fitting model for FA included log(distance), WMH type (tract-WMH and nearby WMH), and their interaction as fixed effects, and random intercept and slope for both participant and tract. For MD, the same model, but without the interaction effect, gave the best fit (lowest Bayesian information criterion). Age and gender effects were tested in a second model. Type III Wald F tests with Kenward-Roger df approximation were used to obtain F and p-values. Due to the small sample of this pilot study, this analysis was not repeated for individual tracts.

## 3. Results

### 3.1 Participant demographics

Data from 52 participants (27 male) with a mean age of 72.2 (standard deviation 0.7) years were used for analysis (Table 1). Out of the 60 participants in the initial sample, Tracula failed to produce viable tractography outputs in six participants, one participant’s WMH were too small to be segmented, while another participant’s WMH did not intersect any of the tracts of interest, and they were therefore excluded from the analysis. Participants varied in Fazekas score and consequently also in number of tracts without presence of WMH. Only 6 participants had less than 8 tracts (out of 18 segmented) with presence of WMH, and most participants (N=46) had at least 8 tracts intersecting WMH. Additionally, the highest percentage of participants had a WMH in the costicospinal tract followed by the bilateral anterior thalamic radiation and right uncinate fasciculus (Table 1). The lowest percentage of participants with a WMH was observed for the bilateral cingulum angular bundle.

**Table 1.** Participant demographics and percentage of WM tracts affected by WMH.

| N (male) | 52 (27) | WM tract | % of N with WMH in WM tract |
| --- | --- | --- | --- |
| Mean age (SD) | 72.2 (0.7) | Anterior thalamic radiation L | 84.6 |
| Total Fazekas score | N | Anterior thalamic radiation R | 82.7 |
| 1 | 7 | Cingulate cingulum L | 75.0 |
| 2 | 9 | Cingulate cingulum R | 78.9 |
| 3 | 10 | Cingulum angular bundle L | 46.2 |
| 4 | 9 | Cingulum angular bundle R | 50.0 |
| 5 | 9 | Corticospinal tract L | 94.2 |
| 6 | 8 | Corticospinal tract R | 92.3 |
| Number of tracts without WMH | N | Inferior longitudinal fasciculus L | 78.9 |
| 0 | 9 | Inferior longitudinal fasciculus R | 76.9 |
| 1-5 | 35 | Forceps Major | 67.3 |
| 6-10 | 2 | Forceps Minor | 75.0 |
| 11-15 | 4 | Superior longitudinal fasciculus, parietal L | 76.9 |
| 16-18 | 2 | Superior longitudinal fasciculus, parietal R | 76.9 |
|  |  | Superior longitudinal fasciculus, temporal L | 78.9 |
|  |  | Superior longitudinal fasciculus, temporal R | 84.6 |
|  |  | Uncinate fasciculus L | 76.9 |
|  |  | Uncinate fasciculus R | 82.7 |

WM=white matter, L=left, R=right, WMH=white matter hyperintensity, N=sample size. WMH % of WM tract and % of N with a WMH in a WM tract, exclude participants without a WMH and those that were excluded after data quality assessment.

### 3.2 Global and local WMH and tract-NAWM diffusion

#### 3.2.1 All tracts combined

We used multiple linear regression to assess the effects of WMH damage on the integrity of NAWM. We analysed FA and MD in separate models, see results in Table 2. In Model 1 we looked at the effects of tract-WMH diffusion, tract % WMHvol and Fazekas in the NAWM water diffusion. For tract-NAWM FA, we found significant effects of mean % WMHvol and tract-WMH FA. For tract-NAMW MD, we found a significant effect of tract-WMH MD.

A second model was run for MD and FA separately including the potential confounders of age and gender. For tract-NAMW FA, the confounders had no significant effect and the significant effects of % WMHvol and tract-WMH FA remained unchanged (Table 2). However, for tract-NAWM MD the introduction of age and gender in the model caused both Fazekas and age to show a trend towards significance. The increase in the effect size of Fazekas between Model 1 and Model 2 for MD can be observed in Table 2.

**Table 2.** Results of the multiple linear regression with local and global damage predictors for tract-NAWM diffusion.

| **Model 1** | **Predictor** | **B<sub>std</sub>** | **F<sub>1,48</sub>** | **p** |
| --- | --- | --- | --- | --- |
| Tract-NAWM FA | Tract-WMH FA | 0.301 | 8.300 | 0.006<sup>a</sup> |
|  | % WMHvol | -0.424 | 8.463 | 0.005<sup>a</sup> |
|  | Fazekas | -0.199 | 1.744 | 0.193 |
| Tract-NAWM MD | Tract-WMH MD | 0.458 | 16.14 | < 0.001<sup>a</sup> |
|  | % WMHvol | 0.194 | 1.370 | 0.248 |
|  | Fazekas | 0.186 | 1.271 | 0.265 |
| **Model 2** | **Predictor** | **B<sub>std</sub>** | **F<sub>1,46</sub>** | **p** |
| Tract-NAWM FA | Tract-WMH FA | 0.309 | 8.387 | 0.006<sup>a</sup> |
|  | % WMHvol | -0.460 | 9.180 | 0.004<sup>a</sup> |
|  | Fazekas | -0.079 | 0.165 | 0.687 |
|  | Age | -0.132 | 0.923 | 0.342 |
|  | Gender (F) | -0.019 | 0.009 | 0.925 |
| Tract-NAWM MD | Tract-WMH MD | 0.449 | 15.84 | < 0.001<sup>a</sup> |
|  | % WMHvol | 0.125 | 0.553 | 0.461 |
|  | Fazekas | 0.412 | 3.864 | 0.055 |
|  | Age | -0.261 | 2.985 | 0.091 |
|  | Gender (F) | -0.003 | < 0.001 | 0.989 |

FA = fractional anisotropy, MD = mean diffusivity, WMH =white matter hyperintensity, NAWM=normal-appearing white matter, B<sub>std</sub> = standardised coefficient
<sup>a</sup> = significant effect

As total Fazekas scores and the tract %WMHvol are correlated (r = 0.744, t = 7.863, p < 0.001; the more the global damage the larger areas of WMH are crossed by the tracts of interest), we checked for collinearity effects in the models. We found variation inflation factors (VIF) of up to 2.4 for %WMHvol and 3.9 for Fazekas, which indicates potential collinearity effects in our models.

We also investigated further the effect of age and Fazekas on water diffusion. MD is known to generally increase with advancing age (Hsu et al., 2010); however, the introduction of age and gender in our MD model indicated a trend of significance for age in the opposite direction to that expected, as well as increasing the effect of Fazekas scores in tract-NAWM MD (with the standardised coefficient of Fazekas increasing from 0.186 to 0.412). We plotted tract-NAWM FA and MD against age for each Fazekas group, and against Fazekas, see Figure 3. The plots showed that although the general trend is that tract-NAWM FA decreased, and MD increased with age (black regression line in Figures 3a and 3c) and Fazekas (blue regression line in Figures 3b and 3d), the trends with age were reversed when the data was divided by Fazekas group (colour regression lines in Figures 3a and 3c), particularly for MD. This indicates the occurrence of a reversal, or “Simpsons”, paradox in the data (Julious and Mullee, 1994), and explains the near significant trends observed in MD Model 2. These figures also show that although participants with higher Fazekas tend to be older, the range of ages overlaps across Fazekas groups. A further analysis of water diffusion changes per Fazekas score can be found in Supplement S1.

**Figure 3.**
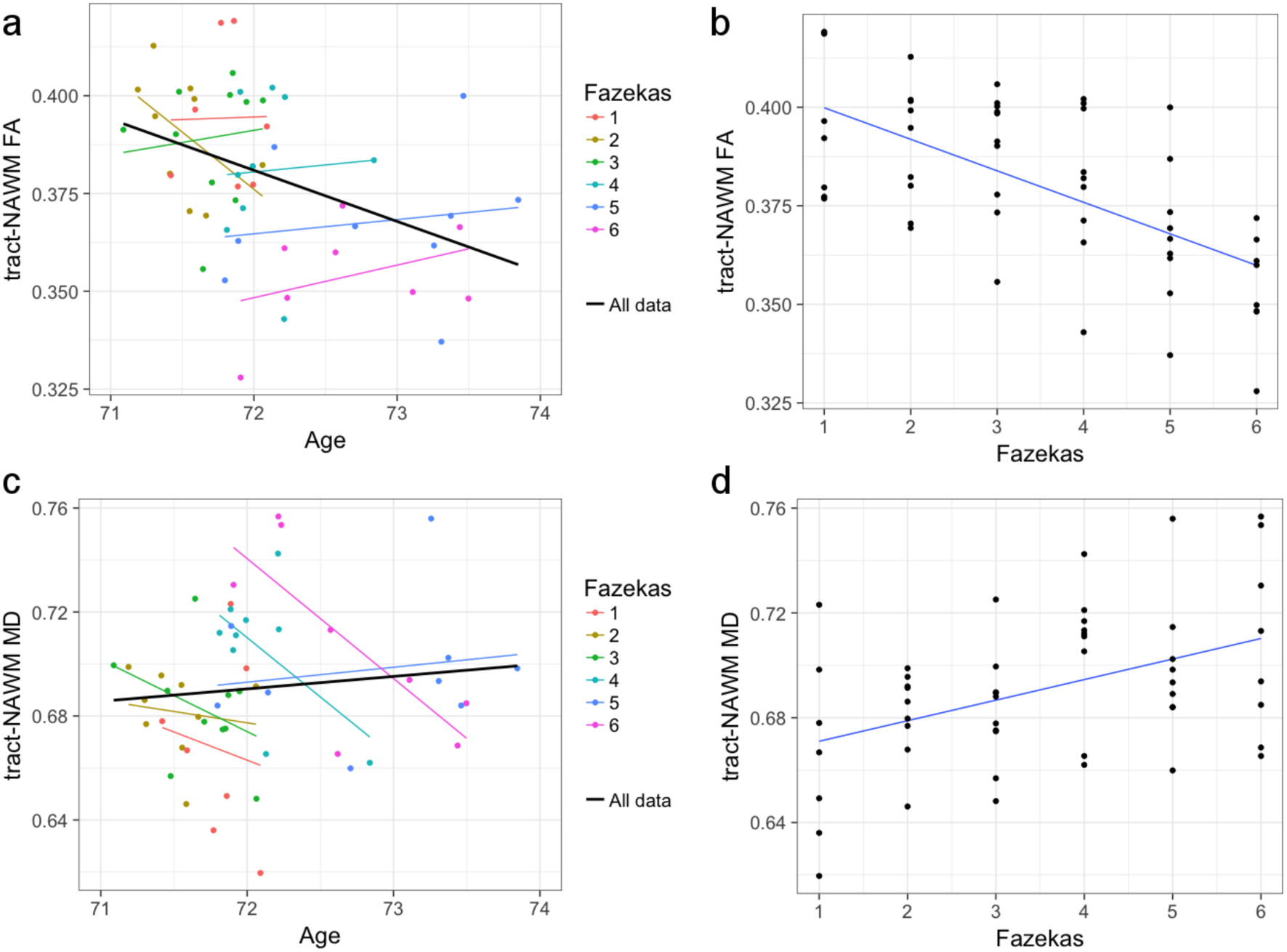
Plot of tract-NAWM water diffusion parameters for all 18 tracts combined. The plots of FA (a) and MD (c) versus age show the overall trend of the parameters with age (black regression line), and the trend for each Fazekas group (colour regression lines). Both parameters present a Simpsons paradox, where the relationship of the parameters with age reverses when Fazekas is considered. (b) and (d) show the relationship of tract-NAMW FA and MD with Fazekas scores. A more comprehensive analysis of diffusion parameters and Fazekas is in Supplement S1.

#### 3.2.2 Individual WM tracts

Median tract-WMH volumes ranged from 0.2 to 2.8% of total WM tract volume, with maximum overlaps seen up to 47% (see Figure 4 and Supplementary Table S2.1).

**Figure 4.**
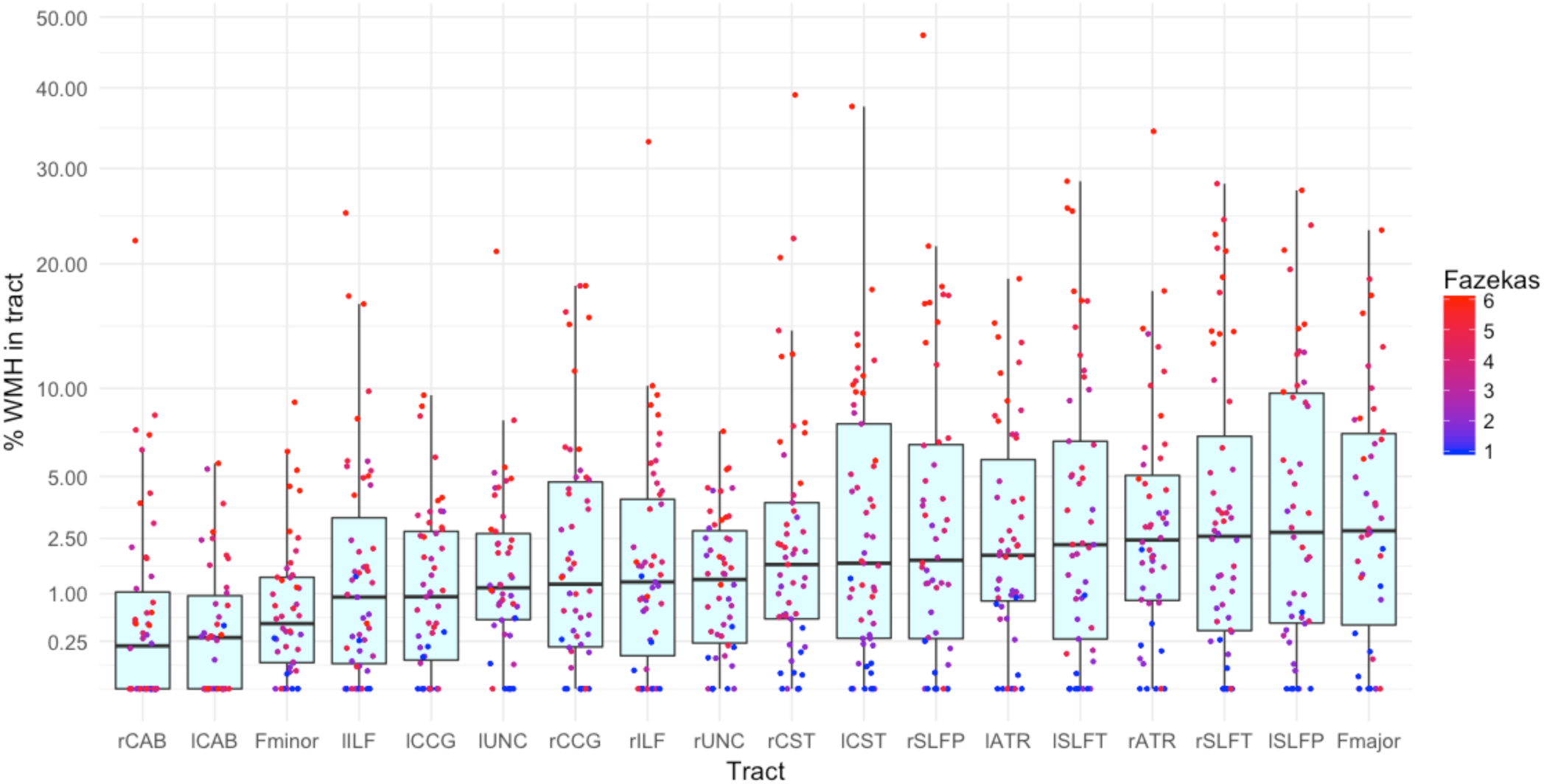
Percentage of the tract intersecting a WMH. Vertical axis is in square-root scale for easier visualisation. Tracts are ordered by median % WMH overlap in tract (less affected on the left, more affected on the right of plot). The boxes represent the lower and upper quartiles and the median measurement (thick line) for each tract, while whiskers indicate the sample minimum and maximum, excluding outliers that differ from the lower and upper quartiles by more than 1.5 times the interquartile range. The dots represent individual data, colour coded by total Fazekas score (1=low, 6=high WMH load). L= left, r =right, Fmajor= forceps major, Fminor= forceps minor, ATR = anterior thalamic radiation, CCG = cingulate cingulum, CAB = cingulum angular bundle, CST = corticospinal tract, ILF = inferior longitudinal fasciculus, SLFp = parietal superior longitudinal fasciculus, SLFt = temporal superior longitudinal fasciculus UNC = uncinate fasciculus.

To assess the predictive effects of WMH damage on the integrity of NAWM in each individual tract we used Model 1 only, which included the effects of tract-WMH diffusion, and tract %WMHvol and Fazekas in the NAWM water diffusion measurements. Full results are shown in (Supplement S2.2).

These data show that tract-WMH FA predicted tract-NAWM FA in the left CAB and right Unc. Tract-WMH MD predicted tract-NAWM MD in the forceps minor, left ATR, left CST, right ATR, right SLFt and right Unc. Tract % WMHvol significantly predicted tract-NAWM FA in the left CST, right CST, right CAB, and tract-NAWM MD in the left Unc. And Fazekas score significantly predicted tract-NAWM FA in the left ILF. The VIF of total Fazekas scores and tract %WMHvol were > 2 for the left and right SLFt and SLFp only, with a maximum VIF of 3.2.

### 3.3 Water diffusion in tract-WMH and tract-NAWM

#### 3.3.1 All tracts

The median FA for all tracts was 0.290 for tract-WMH and 0.380 for tract-NAWM; the median MD for all tracts was 0.895 × 10^-3^mm^2^/s for tract-WMH and 0.690 × 10^-3^mm^2^/s tract-NAWM.

Model 1 showed that tract-WMH had reduced FA (estimate = −0.082, F(1, 23.6) = 82.10, p < 0.001) and increased MD (estimate = 0.197 × 10^-3^ mm^2^/s; F(1, 41.3) = 207.6, p < 0.001) in comparison with tract-NAWM. Model 2, including age and gender as covariates, showed a significant effect of age in FA, with lower FA for older participants (estimate = −0.126, F (1, 49.3) = 8.06, p = 0.007).

#### 3.3.2. Individual tracts

Tract-WMH, in comparison with tract-NAWM, showed decreased FA (p < 0.01) in all tracts except for the left and right cingulum angular bundle, and increased MD (p < 0.01) in all tracts (Supplementary section S2.3).

### 3.4 Water diffusion in tract-WMH and tract-NAWM spatial contours

#### 3.4.1 All tracts

We observed some outliers in the model caused by CSF contamination in some NAWM contours. Contours with MD > 10^-3^mm^2^/s were therefore excluded from analysis as we found MD unlikely to be over this value in white matter in this population (Muñoz Maniega et al., 2015). Only three data points were excluded, two 2mm contours and one 4mm contours, all from the forceps minor.

Model 1 showed that FA increases significantly, (estimate = 0.035, F(1, 21.5) = 78.2, p < 0.001), while MD decreases significantly (estimate = −0.085, F(1, 36.9) = 202.0, p < 0.001), and logarithmically with the distance from the WMH. The top row of Table 3 shows the median water diffusion values obtain at each distance, while Figure 5 shows (in light colour) the boxplots for each distance to tract-WMH for all tracts combined. Figure 6 shows the model predicted water diffusion changes in tract-NAWM with distance from tract-WMH.

**Figure 6.**
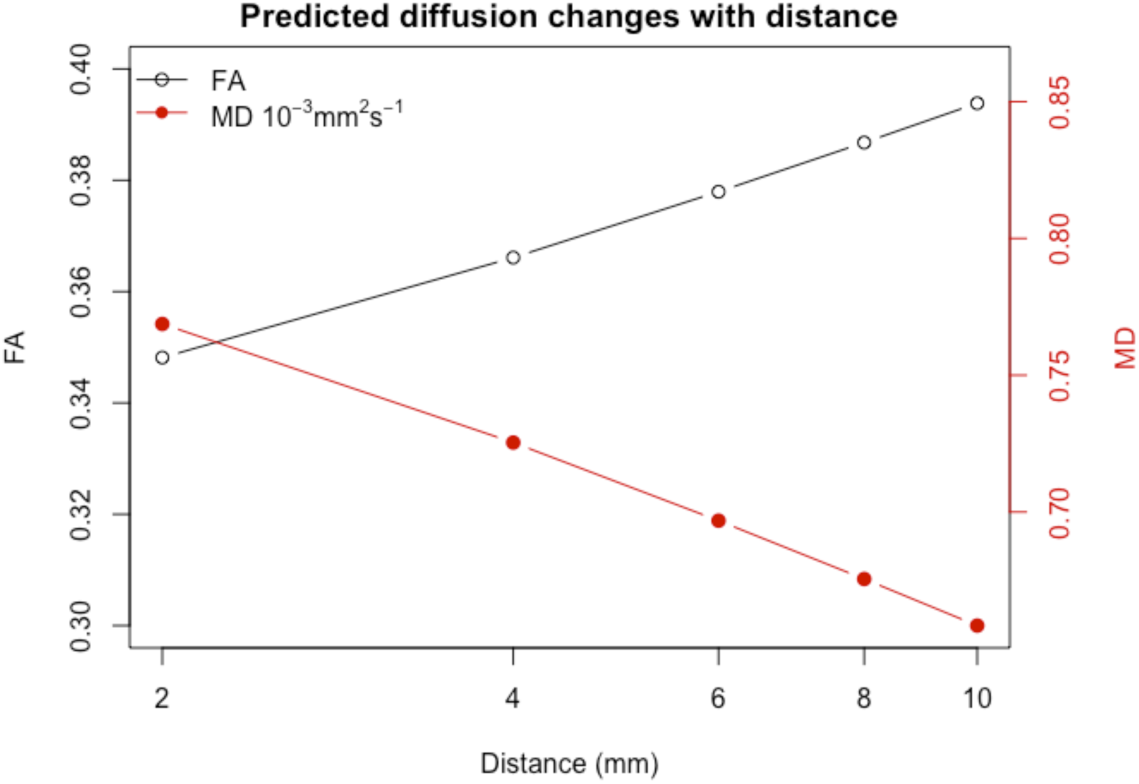
Model predicted water diffusion changes in tract-NAWM with distance from tract-WMH, FA in black, MD in red. Distance scale is logarithmic for easier visualisation

Age and gender had no significant effect in Model 2.

**Table 3.**
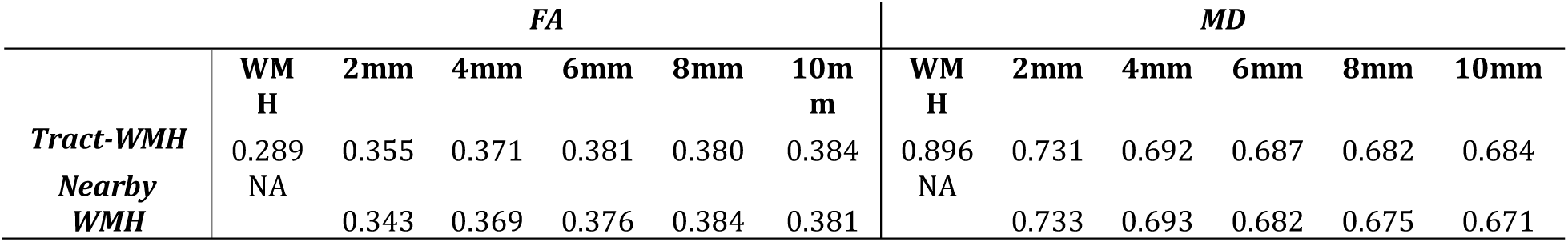
Median diffusion values (FA and MD (× 10-3mm2/s)) of tract-WMH and tract-NAWM spatial distances, for all WM tracts combined and each WMH type. Distances are displayed in mm away from the WMH.

|  | FA |  |  |  |  |  | MD |  |  |  |  |  |
| --- | --- | --- | --- | --- | --- | --- | --- | --- | --- | --- | --- | --- |
|  | WM<br>H | 2mm | 4mm | 6mm | 8mm | 10m<br>m | WM<br>H | 2mm | 4mm | 6mm | 8mm | 10mm |
| **Tract-WMH** | 0.289 | 0.355 | 0.371 | 0.381 | 0.380 | 0.384 | 0.896 | 0.731 | 0.692 | 0.687 | 0.682 | 0.684 |
| **Nearby**<br>**WMH** | NA | 0.343 | 0.369 | 0.376 | 0.384 | 0.381 | NA | 0.733 | 0.693 | 0.682 | 0.675 | 0.671 |

#### 3.4.2 Individual tracts

A linear mixed model with log(distance+1) as a fixed effect and random intercept and slope per study participants was used for each tract. The models showed that MD changes significantly with distance for all tracts while FA changes significantly with distance all tracts except R and L CAB. Supplement S2.4 shows plots and overall F-test results.

### 3.5 Water diffusion in tract-NAWM spatial contours: tract-WMH and nearby-WMH compared

Figure 5 shows the changes of water diffusion in tract-NAWM with distance to the tract-WMH (intersecting) and nearby WMH. Table 3 shows the median values of FA and MD at distances from each of the WMH type.

For FA, the model showed significant effects for WMH type (estimate = −0.028, F (1, 7288.7) = 30.5, p < 0.001), with higher FA for tract-WMH contours than for nearby-WMH contours, and for the interaction of log(Distance):WMH type (estimate = 0.014, F (1, 7292.9) = 23.6, p < 0.001). The main effect of distance was not significant (estimate = 0.011, F (1, 25.5) = 2.74, p = 0.110).

For MD, WMH type had a significant effect (estimate = −0.006, F (1, 7281.9) = 20.1, p < 0.001), with higher MD for tract-WMH contours than for nearby-WMH contours. The log(distance) effect was also significant (estimate = −0.032, F (1,33.2) = 111.6, p < 0.001).

Model 2 show no significant effects of age or gender in either FA or MD.

## 4. Discussion

In this study we looked at water diffusion parameters of WMH and NAWM, focusing specifically on the major brain WM tracts. We described quantitatively the patterns of overlap between these tracts and WMH, and how these are related to global WMH burden. We also observed that water diffusion values in tract-NAWM were predicted by the values in tract-WMH. FA in tract-NAWM was also predicted by the percentage of tract-WMH volume, while MD in tract-NAWM was predicted—albeit as a trend—by overall WMH burden. We detected that water diffusion values were abnormal not only in the WMH-tract segment, but importantly, that the abnormalities continued within the NAWM of the tract. We described quantitively the changes in tract-NAWM with distance from the tract-WMH, with lessening abnormalities as we move further away and along the WM tract. A similar distance pattern of abnormalities in tract-NAWM water diffusion metrics was observed for nearby-WMH. This indicates that tissue microstructural changes in WM tracts spread beyond the visible damage of the WM, and also suggests that these NAWM changes are associated with both WMH crossed by the tract, as well as those nearby.

We observed expected microstructural tract-WMH abnormalities, following the pattern of higher MD and lower FA, for the majority of the 18 WM tracts assessed. Such microstructural abnormalities may be caused by a variety of microvascular dysfunctions, resulting in interstitial oedema, inflammation, ischaemia and damage to the myelin sheet of WM tracts, ultimately leading to visible WMH (Wardlaw et al., 2017a). The cingulum angular bundle was one exception where FA in WMH-tract was not abnormal, which might be explained by an underpowered analysis as only ∼50% of the sample had a WMH in the cingulum angular bundle.

To our knowledge, only a few studies of brain ageing have specifically analysed WM tracts intersecting with WMH (Langen et al., 2017; Maillard et al., 2015; Seiler et al., 2018; Taylor et al., 2017). They generally found the largest rates of overlap in the anterior thalamic radiation, inferior longitudinal fasciculus, forceps major, posterior thalamic radiation and the inferior fronto-occipital fasciculus. We found similar rates of overlap for the forceps major and anterior thalamic radiation in our study, but also observed large overlap in other tracts, such as in the parietal and temporal superior longitudinal fasciculi (median 2.0-2.7%, with several subjects with > 25%, Figure 4). Differences in the patterns of overlap with previous studies could be caused by different tractography methods employed, or by the different age ranges of the samples, as the prevalence of WMH is highly dependent on age (de Leeuw et al., 2001).

In the current study, the percentage of participants with a WMH was highest in the corticospinal tract (>92%), with a median WMH volume of 1.7%. This tract was not included in previous studies. The high incidence may be explained by the proximity of this tract to the lateral ventricles, and the common occurrence of periventricular WMH (Valdés Hernández et al., 2015). Similarly, the percentages of participants with a WMH in the forceps minor and the uncinate fasciculus, both tracts prominently passing through periventricular areas, were high (75-82.7%), but with smaller WMH volumes (median 0.5-1.3% WMH), potentially due to the typical pattern of distribution of WMH. The patterns of overlap that we observe do, however, agree with previous studies on the distribution of WMH (Habes et al., 2016; Lambert et al., 2016). Additionally, we observed that the cingulum angular bundle was least often affected by WMH (46.2-50%). This was reported previously (Seiler et al., 2018), and might be explained by the smaller size of this tract, or because its section within the periventricular area is mostly located medially of the posterior part of the lateral ventricle, whereas periventricular WMH area mainly observed laterally and near the anterior or posterior ends of the lateral ventricle (Valdés Hernández et al., 2015). It can also be noted from Figure 4, that most participants with a low total Fazekas score did not have WMH overlapping with the cingulum angular bundle, while other tracts do overlap with WMH for a wider range of Fazekas scores. As expected, the distribution of Fazekas score in Figure 4 shows that those with lower Fazekas scores tend to have less overlap with all tracts.

Previous studies have observed that the ratio of overlap between WMH and individual tracts predicted the mean FA in the whole-tract, independent of overall brain WMH burden (Maillard et al., 2015; Seiler et al., 2018). This effect could have been biased due to inclusion of WMH in the measurement of whole-tract FA, i.e. the larger proportion of WMH in a tract, the lower the whole-tract FA. However, in our study, we observed that higher tract-WMH volume specifically predicts worse FA abnormalities in the tract-NAWM, also independently of overall WMH burden (Fazekas score), indicating that tract-WMH has a genuine effect on tract-NAWM FA. Those studies did not report the effect of WMH overlap on tract MD. We found that tract-WMH volume did not predict MD abnormalities in tract-NAWM, but instead, overall WMH burden had a predictive trend of the level of MD in tract-NAWM. This suggests that tract-NAWM changes in water content and mobility, reflected by MD abnormalities, could be more related to diffuse WM damage typically seen in the ageing brain, whereas tract-NAWM changes in myelin and axonal packing, reflected by FA, are more related to tract-specific localised WM damage or to premorbid WM integrity with lower FA predisposing to development of WMH. These results, however, need to be interpreted with caution due to the potential collinearity effects between %WMHvol and Fazekas in the models.

We also found a Simpson’s paradox between Fazekas and age in tract-NAWM. Figure 3 shows the expected general trend of increasing MD and decreasing FA with both increasing Fazekas and age; however, the expected trend of the water diffusion parameters with age is reversed within each Fazekas group. An explanation of the age effect reversal within a Fazekas group could be that an older person with the same amount of WM damage as a younger person (same Fazekas group) could be generally in better health (hence developed the same damage at an older age, even within the very narrow age range in the LBC1936), and their health could be reflected in their baseline WM integrity. A similar paradoxical effect was recently observed in a minor stroke study, where patients who were older at the time of stroke had generally better premorbid intelligence (NART), which could also be reflecting better health and hence delay in stroke onset (Makin et al., 2018). Lower childhood IQ has been associated with higher WMH load (Backhouse et al., 2017), and worse white matter structural integrity (FA) (Deary et al., 2006). Better baseline WM integrity could therefore play a role in delaying WM damage either directly, or mediated by premorbid cognition, which could influence later health. Future work should investigate the combined effects of WM integrity, age, and premorbid cognition in the accumulation of brain WMH.

In addition, both tract-NAWM FA and MD were predicted by tract-WMH FA and MD respectively, suggesting that microstructural changes in tract-NAWM are not only associated with overall WMH burden (for MD) or tract-specific WMH burden (for FA), but also with the severity of the microstructural changes in tract-WMH. These predictive patterns were slightly supported by our exploratory analysis assessing them in individual tracts where we observed a similar prediction pattern for several tracts.

The fact that overall microstructural integrity of tract-NAWM become more abnormal as WMH burden increases was further supported by the specific analysis of changes in diffusion with Fazekas scores, which showed that diffusion of tract-NAWM deteriorates further for higher Fazekas groups (Supplement S1). This corroborates similar effects we previously found in the study of whole-brain WM in stroke patients (Muñoz Maniega et al., 2017) and healthy elderly (Muñoz Maniega et al., 2015). There have been other studies reporting similar results (Firbank et al., 2003; Pelletier et al., 2016). This, together with the predictive effects of tract-WMH water diffusion metrics on tract-NAWM diffusion values, suggest that the microstructural changes observed in both WMH and NAWM are part of the same overall pathological process.

A distance gradient of microstructural abnormalities was observed in tract-NAWM. We observed that FA and MD in tract-NAWM are abnormal at 2-10 mm away from the tract-WMH visible edge. Both parameters change consistently in a gradual manner, with FA decreasing and MD increasing as we get closer to the visible lesion. These results are not entirely in line with previous studies conducted by our group (Muñoz Maniega et al., 2015; Wardlaw et al., 2017b). In these studies, we *qualitatively* observed a similar distance pattern for whole-brain NAWM up to and including 4 mm distance for FA and 8 mm for MD, but at further distances the abnormalities in the NAWM apparently increased. This might be explained by the fact that in our previous work we looked at whole-brain NAWM and were not able to exclude the location bias due to WMH located in areas of high baseline FA (low baseline MD). Spatial contours further away from the lesions would encircle lower baseline FA areas (higher baseline MD), and thus would induce a so-called increase of abnormalities. The current study did not suffer from a location bias, because water diffusion in the NAWM was measured within the same tract (stable baseline water diffusion).

It has been suggested that, from a biological perspective, we could assume that a WMH in a white matter tract would influence the tissue integrity along the rest of the tract more strongly than in the rest of the (non-directly connected) WM (Maillard et al., 2011). However, our data shows for the first time that water diffusion within tract-NAWM displays similar patterns for both intersecting and nearby WMH. FA was slightly lower in nearby WMH contours of NAWM in relation to tract-WMH contours (although the significant interaction term between WMH type and distance indicates a cross-over interaction in some contours) while the MD was slightly higher for tract-WMH contours. However, the estimated differences were small for both parameters, and there was a comparable gradual decrease of abnormalities as distance from the edge of both types of WMH increased (Figure 5). This may suggest that age-related WM damage does not propagate predominantly along tract axons, but propagates via different mechanisms, regardless of whether the WMH is in the tract or next to it. For example, age-related WMH are associated with other small vessel disease (SVD) features such as widening of perivascular spaces (Fazekas et al., 1993; Munoz et al., 1993; Schmidt et al., 2011), that reflect microvascular dysfunction including abnormal blood-brain barrier leakage, impaired vasoreactivity and impaired pulsatility. Our results, together with the finding in other SVD populations that MD is more abnormal than FA measures in NAWM adjacent to WMH (Maclullich et al., 2009; Muñoz Maniega et al., 2017) support the relevance of other potential channels for the propagation of tissue damage, independently of the direct connections of WM tracts.

Water diffusion abnormalities within the NAWM are suggestive of tissue damage not yet visible on a lesion level. These water diffusion abnormalities are a precursor to lesion extension or development within this area of NAWM (de Groot et al., 2013; Maillard et al., 2014, 2013; Mayo et al., 2017), which reflect the general association between higher WMH load predicting worse NAWM MD and more WMH growth. As lesions are associated with cognitive abnormalities (Biesbroek, 2016; Reginold et al., 2015; Valdés Hernández et al., 2013a), such lesion development in NAWM areas suggests that pre-lesional NAWM water diffusion abnormalities may already play a role in cognitive functioning (Baykara et al., 2016; Jokinen et al., 2013). Previous studies have found that the rate of WMH growth was heterogeneous, occurring more rapidly within some association and projection tracts (Lambert et al., 2016). A longitudinal study of this growth at the tract level, and with respect to the location of tract-WMH and nearby WMH induced changes in the tract-NAWM, could help explain the patterns of WMH progression. Our future efforts are aimed at elucidating the patterns of lesion formation following water diffusion abnormalities in NAWM, as well as at clarifying the associations between cognitive dysfunctioning and diffusion abnormalities in both tract-WMH and tract-NAWM.

This study has some limitations. Firstly, it has a relatively small sample, mainly for the individual tract comparisons where there were less cases with WMH in particular tracts. However, the current study was conducted as a pilot study and its results will be used as motivation for investigating WMH-tract interactions in the whole LBC1936 sample. The current study can be further extended to focus on relating water diffusion in tract-WMH and tract-NAWM to premorbid cognition, current cognitive functioning, and on the development of water diffusion abnormalities in tract-WMH and tract-NAWM over time. Secondly, the closer spatial contours were not available for every tract as some nearby WMH were too far away for a 2-or 4-mm contour to cross with the tract. This means that there was less data contributing to the calculation of water diffusion for 2 and 4 mm and their measurements might be less accurate than for the tract-WMH contours, the patterns of change with distance will therefore need to be corroborated in a larger sample.

In conclusion, tract-WMH showed WM microstructural changes suggestive of tissue damage, that were predictive of similar—albeit of a lesser degree—changes in tract-NAWM. Microstructural changes in tract-NAWM became less pronounced along the tract and further away from the tract-WMH, with a comparable distance pattern away from the nearby (not intersecting) WMH. Overall, these results suggest tissue damage in WM tracts beyond visible lesions, which may contribute to cortical disconnection and changes in cognitive functioning. Our future efforts are aimed at elucidating the relationship between microstructural changes of nearby and tract-WMH, tract-NAWM and cognitive functioning at older ages, and at the development of tissue changes in tract-NAWM over time.

**Supplement S1 – Diffusion changes per Fazekas score in tract-WMH and tract-NAWM**

We used repeated measurements linear mixed models to assess between-Fazekas score differences in diffusion for both tract-WMH and tract-NAWM. We used the total Fazekas score as a fixed effect, random intercept for participant and tract and random slope for tract. Data for each tract was considered as a repeat measurement. A second model, including age and gender was used to test the effects of these potential confounders. Pairwise comparisons were performed for Fazekas scores, with a significance threshold set at p<0.01 to take multiple comparisons into consideration.

## Results

FA of tract-NAWM was different between Fazekas groups (F(5,47.1)=5.64, p < 0.001, while there was a trend for tract-NAWM MD (F(5,47.9)=2.50, p=0.044) (Table S1; Figure S1), showing an overall pattern of decreased FA and increased MD with increasing Fazekas score. Pairwise analysis showed that tract-NAWM FA was significantly higher for Fazekas 1, 2, 3 and 4 compared with Fazekas 6, and Fazekas 1 and 3 compared to Fazekas 5. Tract-NAWM MD was significantly lower in Fazekas 1 compared with Fazekas 4 and 6. There was no significant effects of age and gender, or significant differences between Fazekas groups for WMH, however the sample size was small, with only 7-10 of participants per Fazekas group, and a lower number of tracts contributing to WMH data for the lower Fazekas groups.

**Table S1.** Median FA and MD (x 10-3mm2/s) for tract-NAWM and tract-WMH per Fazekas scores. Significance values are uncorrected for multiple comparisons.

|  | **Tract-NAWM** |  | **Tract-WMH** |  |
| --- | --- | --- | --- | --- |
| **Fazekas score** | FA | MD | FA | MD |
| **1** | 0.392<sup>a(p=0.002),b(p=0.008)</sup> | 0.667 | 0.335 | 0.869 |
| **2** | 0.395<sup>a(p=0.002)</sup> | 0.686 | 0.289 | 0.932 |
| **3** | 0.395<sup>a(p=0.001),b(p=0.008)</sup> | 0.683 | 0.308 | 0.850 |
| **4** | 0.382<sup>a(p=0.003)</sup> | 0.712<sup>c(p=0.008)</sup> | 0.293 | 0.926 |
| **5** | 0.367 | 0.693 | 0.282 | 0.933 |
| **6** | 0.355 | 0.704<sup>c(p=0.006)</sup> | 0.273 | 0.910 |
| FA = fractional anisotropy, MD = mean diffusivity, NAWM = normal-appearing white matter, WMH = white matter hyperintensity.<br><sup>a</sup> = significant difference in comparison with Fazekas 6<br><sup>b</sup> = significant difference in comparison with Fazekas 5<br><sup>c</sup> = significant difference in comparison with Fazekas 1 |  |  |  |  |

**Table S2.1.** Descriptive statistics of the percentage of tract volume intersecting WMH

| % WMH | L/R | Mean | Median | Min | Max | Q1 | Q3 | IQR |
| --- | --- | --- | --- | --- | --- | --- | --- | --- |
| **F Major** | n/a | 5.0 | 2.8 | 0.0 | 23.3 | 0.5 | 7.2 | 6.7 |
| **F Minor** | n/a | 1.2 | 0.5 | 0.0 | 9.1 | 0.1 | 1.4 | 1.3 |
| **ATR** | L | 3.8 | 2.0 | 0.0 | 18.6 | 0.9 | 5.9 | 5.0 |
| **ATR** | R | 4.4 | 2.5 | 0.0 | 34.4 | 0.9 | 5.1 | 4.2 |
| **CCG** | L | 1.8 | 0.9 | 0.0 | 9.5 | 0.1 | 2.7 | 2.7 |
| **CCG** | R | 3.4 | 1.2 | 0.0 | 18.0 | 0.2 | 4.7 | 4.5 |
| **CAB** | L | 0.8 | 0.3 | 0.0 | 5.6 | 0.0 | 1.0 | 1.0 |
| **CAB** | R | 1.5 | 0.2 | 0.0 | 22.2 | 0.0 | 1.0 | 1.0 |
| **CST** | L | 4.6 | 1.7 | 0.0 | 37.6 | 0.3 | 7.8 | 7.5 |
| **CST** | R | 4.0 | 1.7 | 0.0 | 39.1 | 0.5 | 3.8 | 3.3 |
| **ILF** | L | 2.8 | 0.9 | 0.0 | 25.1 | 0.1 | 3.3 | 3.2 |
| **ILF** | R | 2.9 | 1.3 | 0.0 | 33.2 | 0.1 | 4.0 | 3.8 |
| **SLFp** | L | 5.8 | 2.7 | 0.0 | 27.5 | 0.5 | 9.7 | 9.2 |
| **SLFp** | R | 5.6 | 1.8 | 0.0 | 47.3 | 0.3 | 6.6 | 6.3 |
| **SLFt** | L | 5.7 | 2.6 | 0.0 | 28.3 | 0.4 | 7.1 | 6.7 |
| **SLFt** | R | 5.5 | 2.3 | 0.0 | 28.5 | 0.3 | 6.8 | 6.5 |
| **UNC** | L | 2.2 | 1.1 | 0.0 | 21.2 | 0.5 | 2.7 | 2.1 |
| **UNC** | R | 1.7 | 1.3 | 0.0 | 7.3 | 0.2 | 2.8 | 2.5 |

WMH =white matter hyperintensity, L= left, R=right, F= forceps, ATR = anterior thalamic radiation, CCG = cingulate cingulum, CAB = cingulum angular bundle, CST = corticospinal tract, ILF = inferior longitudinal fasciculus, SLFp = parietal superior longitudinal fasciculus, SLFt = temporal superior longitudinal fasciculus UNC = uncinate fasciculus. Q1 = first quartile, Q3 = third quartile, IQR = interquartile range. Note: 'Max' values are inclusive of outliers.

**Table S2.3.** Means and paired t-test results of diffusion in tract-NAWM and tract-WMH (N = 52)

| *tract* | *tract-NAWM FA* | *tract-WMH FA* | *t* | *est diff* | *L conf. int* | *H conf. int* | *p* |
| --- | --- | --- | --- | --- | --- | --- | --- |
| ***F Major*** | 0.454 | 0.347 | 6.852 | 0.105 | 0.074 | 0.137 | < 0.001 |
| ***F Minor*** | 0.375 | 0.244 | 8.325 | 0.129 | 0.098 | 0.16 | < 0.001 |
| ***L ATR*** | 0.347 | 0.253 | 12.11 | 0.092 | 0.077 | 0.107 | < 0.001 |
| ***L CCG*** | 0.423 | 0.272 | 12.49 | 0.148 | 0.124 | 0.172 | < 0.001 |
| ***L CAB*** | 0.308 | 0.303 | 0.078 | 0.002 | -0.051 | 0.055 | 0.938 |
| ***L CST*** | 0.476 | 0.404 | 5.080 | 0.070 | 0.042 | 0.098 | < 0.001 |
| ***L ILF*** | 0.385 | 0.316 | 5.192 | 0.065 | 0.040 | 0.091 | < 0.001 |
| ***L SLFp*** | 0.361 | 0.298 | 5.473 | 0.059 | 0.037 | 0.08 | < 0.001 |
| ***L SLFt*** | 0.388 | 0.308 | 8.793 | 0.076 | 0.058 | 0.093 | < 0.001 |
| ***L UNC*** | 0.34 | 0.239 | 14.84 | 0.100 | 0.086 | 0.114 | < 0.001 |
| ***R ATR*** | 0.347 | 0.254 | 10.97 | 0.091 | 0.074 | 0.108 | < 0.001 |
| ***R CCG*** | 0.403 | 0.272 | 15.71 | 0.128 | 0.111 | 0.144 | < 0.001 |
| ***R CAB*** | 0.309 | 0.269 | 1.751 | 0.044 | -0.008 | 0.096 | 0.092 |
| ***R CST*** | 0.459 | 0.379 | 6.257 | 0.078 | 0.053 | 0.103 | < 0.001 |
| ***R ILF*** | 0.38 | 0.33 | 4.567 | 0.042 | 0.023 | 0.061 | < 0.001 |
| ***R SLFp*** | 0.368 | 0.283 | 9.931 | 0.082 | 0.065 | 0.099 | < 0.001 |
| ***R SLFt*** | 0.379 | 0.298 | 8.832 | 0.080 | 0.062 | 0.098 | < 0.001 |
| ***R UNC*** | 0.329 | 0.228 | 10.29 | 0.097 | 0.078 | 0.117 | < 0.001 |
| *tract* | *tract-NAWM MD* | *tract-WMH MD* | *t* | *est diff* | *L conf. int* | *H conf. int* | *p* |
| ***F Major*** | 0.7 | 0.974 | -14.01 | -0.273 | -0.312 | -0.233 | < 0.001 |
| ***F Minor*** | 0.77 | 0.968 | -9.892 | -0.193 | -0.233 | -0.154 | < 0.001 |
| ***L ATR*** | 0.706 | 0.959 | -15.56 | -0.247 | -0.279 | -0.215 | < 0.001 |
| ***L CCG*** | 0.648 | 0.863 | -10.71 | -0.210 | -0.250 | -0.171 | < 0.001 |
| ***L CAB*** | 0.705 | 0.858 | -5.045 | -0.146 | -0.206 | -0.086 | < 0.001 |
| ***L CST*** | 0.623 | 0.739 | -9.520 | -0.114 | -0.138 | -0.090 | < 0.001 |
| ***L ILF*** | 0.734 | 0.977 | -11.24 | -0.238 | -0.281 | -0.196 | < 0.001 |
| ***L SLFp*** | 0.673 | 0.867 | -12.47 | -0.190 | -0.221 | -0.160 | < 0.001 |
| ***L SLFt*** | 0.67 | 0.878 | -13.72 | -0.203 | -0.233 | -0.173 | < 0.001 |
| ***L UNC*** | 0.743 | 0.991 | -14.86 | -0.246 | -0.279 | -0.212 | < 0.001 |
| ***R ATR*** | 0.707 | 0.923 | -13.33 | -0.210 | -0.242 | -0.178 | < 0.001 |
| ***R CCG*** | 0.638 | 0.862 | -13.81 | -0.218 | -0.250 | -0.186 | < 0.001 |
| ***R CAB*** | 0.703 | 0.822 | -3.474 | -0.112 | -0.179 | -0.046 | 0.002 |
| ***R CST*** | 0.625 | 0.765 | -8.397 | -0.140 | -0.173 | -0.106 | < 0.001 |
| ***R ILF*** | 0.742 | 0.993 | -13.45 | -0.245 | -0.282 | -0.209 | < 0.001 |
| ***R SLFp*** | 0.664 | 0.875 | -13.97 | -0.207 | -0.237 | -0.177 | < 0.001 |
| ***R SLFt*** | 0.659 | 0.856 | -13.79 | -0.193 | -0.221 | -0.165 | < 0.001 |
| ***R UNC*** | 0.746 | 0.956 | -9.736 | -0.205 | -0.248 | -0.163 | < 0.001 |

WM=white matter, L= left, R=right, FA = fractional anisotropy, MD = mean diffusivity in ( $10^{-3}\text{mm}^2/\text{s}$ ), WMH =white matter hyperintensity, NAWM=normal-appearing white matter, F=forceps, ATR = anterior thalamic radiation, CCG = cingulate cingulum, CAB = cingulum angular bundle, CST = corticospinal tract, ILF = inferior longitudinal fasciculus, SLFp = parietal superior longitudinal fasciculus, SLFt = temporal superior longitudinal fasciculus UNC = uncinate fasciculus.

**Table S2.4.** Results of the linear mixed models comparing tract-WMH and the tract-NAWM at 5 distances for each individual tract (N = 52)

| tract | FA |  |  |  |  | MD |  |  |  |  |
| --- | --- | --- | --- | --- | --- | --- | --- | --- | --- | --- |
|  | Estimate | F | Df | Df.res | p | Estimate | F | Df | Df.res | p |
| **F Major** | 0.060 | 52.5 | 1 | 33 | < 0.001 | -0.125 | 233 | 1 | 33 | < 0.001 |
| **F Minor** | 0.063 | 94.1 | 1 | 38 | < 0.001 | -0.085 | 112 | 1 | 38 | < 0.001 |
| **L ATR** | 0.040 | 99.9 | 1 | 43 | < 0.001 | -0.108 | 256 | 1 | 43 | < 0.001 |
| **L CCG** | 0.055 | 109 | 1 | 38 | < 0.001 | -0.086 | 86.9 | 1 | 38 | < 0.001 |
| **L CAB** | 0.001 | 0.02 | 1 | 23 | 0.904 | -0.050 | 20 | 1 | 23 | < 0.001 |
| **L CST** | 0.031 | 21.3 | 1 | 46 | < 0.001 | -0.052 | 100 | 1 | 46 | < 0.001 |
| **L ILF** | 0.026 | 21.4 | 1 | 40 | < 0.001 | -0.104 | 133 | 1 | 40 | < 0.001 |
| **L SLFp** | 0.024 | 24 | 1 | 39 | < 0.001 | -0.085 | 167 | 1 | 39 | < 0.001 |
| **L SLFt** | 0.030 | 48.2 | 1 | 40 | < 0.001 | -0.089 | 230 | 1 | 40 | < 0.001 |
| **L UNC** | 0.034 | 75.2 | 1 | 39 | < 0.001 | -0.101 | 214 | 1 | 39 | < 0.001 |
| **R ATR** | 0.040 | 65 | 1 | 42 | < 0.001 | -0.091 | 175 | 1 | 42 | < 0.001 |
| **R CCG** | 0.052 | 211 | 1 | 40 | < 0.001 | -0.089 | 157 | 1 | 40 | < 0.001 |
| **R CAB** | 0.018 | 1.76 | 1 | 25 | 0.196 | -0.053 | 13.3 | 1 | 25 | 0.001 |
| **R CST** | 0.034 | 34.3 | 1 | 47 | < 0.001 | -0.062 | 86.8 | 1 | 47 | < 0.001 |
| **R ILF** | 0.016 | 9.25 | 1 | 39 | 0.004 | -0.111 | 217 | 1 | 39 | < 0.001 |
| **R SLFp** | 0.036 | 71.5 | 1 | 39 | < 0.001 | -0.091 | 215 | 1 | 39 | < 0.001 |
| **R SLFt** | 0.036 | 70.4 | 1 | 43 | < 0.001 | -0.084 | 219 | 1 | 43 | < 0.001 |
| **R UNC** | 0.041 | 83.2 | 1 | 42 | < 0.001 | -0.088 | 98.8 | 1 | 42 | < 0.001 |

WM=white matter, L= left, R=right, FA = fractional anisotropy, MD = mean diffusivity in ( $10^{-3}\text{mm}^2/\text{s}$ ), F=forceps, ATR = anterior thalamic radiation, CCG = cingulate cingulum, CAB = cingulum angular bundle, CST = corticospinal tract, ILF = inferior longitudinal fasciculus, SLFp = parietal superior longitudinal fasciculus, SLFt = temporal superior longitudinal fasciculus, UNC = uncinate fasciculus.

**Supplement S2 – Analysis of individual WM tracts**

### Section S2.1. Percentage of tract intersecting WMH

### Section S2.2 Multiple regression results for global and local WMH effects on tract-NAWM: individual tracts

The linear regression using Model 1, to check for significant predictive values of individual tract-WMH diffusion, percentage WMH volume and Fazekas scores in individual tract-NAWM diffusion, produced the following significant effects.

Tract-NAWM FA was significantly predicted by tract-WMH FA in the left CAB (B_std_ = −0.585, t(20) = 3.572, p = 0.002) and right UNC (B_std_ = −0.478, t(20) = 3.416, p = 0.001); by mean percentage WMH volume in the left CST (B_std_ = −0.521, t(43) = −3.246 p = 0.002), right CST (B_std_ = −0.531, t(44) = −3.724, p = 0.001), and right CAB (B_std_ = −0.473, t(22) = −3.354, p = 0.003); and by Fazekas score in the left ILF (B_std_ = −0.516, t(37) = −3.447, p = 0.001)

Tract-NAWM MD was significantly predicted by tract-WMH MD in the forceps minor (B_std_ = 0.478, t(35) = 3.204, p = 0.003), left ATR (B_std_ = 0.390, t(40) = 3.012, p = 0.004), left CST (B_std_ = 0.484, t(43) = 3.513, p = 0.001), right ATR (B_std_ = 0.491, t(39) = 4.024, p < 0.001), right SLFt (B_std_ = 0.402, t(40) = 2.897, p = 0.006) and right Unc (B_std_ = 0.378, t(39) = 3.150, p = 0.003); and by mean percentage WMH volume in the left Unc (B_std_ = 0.414, t(36) = 3.145, p = 0.003).

### Section S2.3. T-test results for tract-WMH in comparison with tract-NAWM: individual tracts

### Section S2.4. F-test results for tract-WMH and tract-NAWM spatial contours: individual tracts

Table S2.4 shows the results from the models assessing the effect of distance to the tract-WMH in tract-NAWM for individual tracts. Figure S2 shows the individual plots per participant and per tract for FA and MD measured in tract-WHM and in tract-NAWM at approximately 2, 4, 6, 8 and 10 mm from the tract-WMH visible edge.

## Acknowledgements

This research and LBC1936 phenotype collection were supported by Research Into Ageing and continues as part of The Disconnected Mind project, funded by Age UK, and by the UK Medical Research Council (G0701120, G1001245 and MR/M013111/1). Magnetic Resonance Image acquisition and analyses were conducted at the Brain Research Imaging Centre, Neuroimaging Sciences, University of Edinburgh (www.bric.ed.ac.uk) which is part of SINAPSE (Scottish Imaging Network – A Platform for Scientific Excellence) collaboration (www.sinapse.ac.uk) funded by the Scottish Funding Council and the Chief Scientist Office. This work was supported by the Centre for Cognitive Ageing and Cognitive Epidemiology, funded by the Medical Research Council and the Biotechnology and Biological Sciences Research Council (MR/K026992/1), the Row Fogo Charitable Trust (BRO-D.FID3668413), the European Union Horizon 2020, PHC-03-15, project No 666881, ‘SVDs@Target’, the Fondation Leducq Transatlantic Network of Excellence for the Study of Perivascular Spaces in Small Vessel Disease, ref no. 16 CVD 05, and the Medical Research Council UK Dementia Research Institute at the University of Edinburgh. We thank the Lothian Birth Cohort 1936 participants who took part in this study, the Lothian Birth Cohort 1936 research team members, and radiographers at the Brain Research Imaging Centre.

---

**Figure S1.**
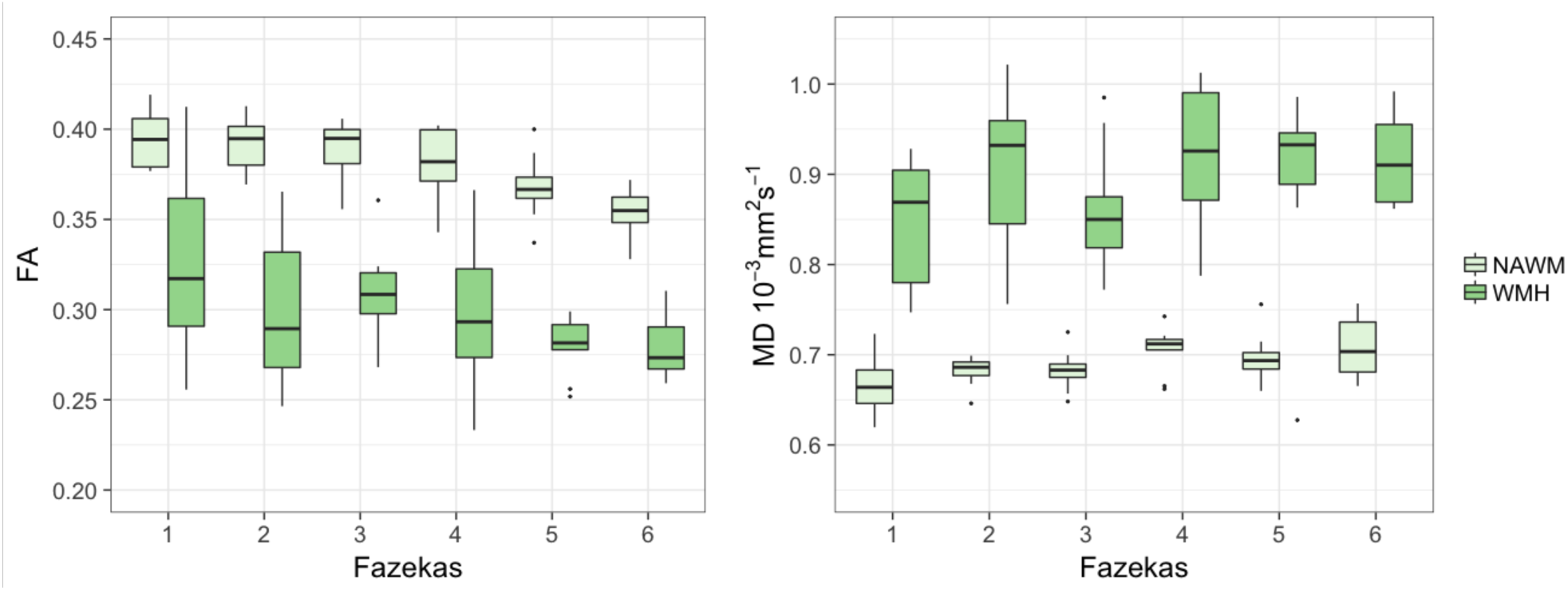
Diffusion values (MD and FA) for tract-WMH and tract-NAWM for all WM tracts combined, ordered by Fazekas.

**Figure S2.**
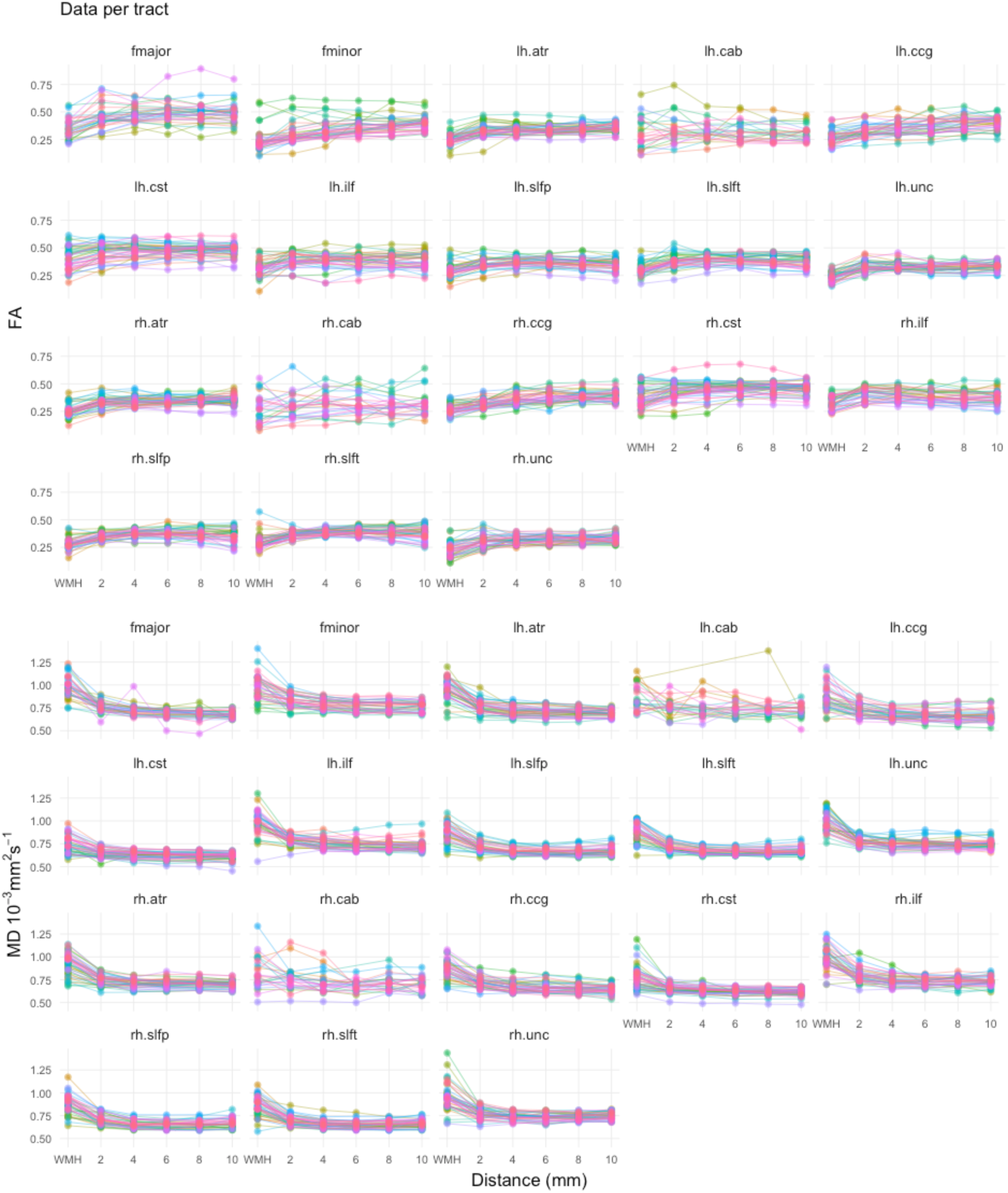
FA (top) and MD (bottom) variation with distance from the visible tract-WMH for each individual tract. Colours represent measurements in individual participants.

